# Enthesis Maturation in Engineered Ligaments is Differentially Driven by Loads that Mimic Slow Growth Elongation and Rapid Cyclic Muscle Movement

**DOI:** 10.1101/2023.03.08.531765

**Authors:** M. Ethan Brown, Jennifer L. Puetzer

## Abstract

Entheses are complex attachments that translate load between elastic-ligaments and stiff-bone via organizational and compositional gradients. Neither natural healing, repair, nor engineered replacements restore these gradients, contributing to high re-tear rates. Previously, we developed a novel culture system which guides ligament fibroblasts in high-density collagen gels to develop early postnatal-like entheses, however further maturation is needed. Mechanical cues, including slow growth elongation and cyclic muscle activity, are critical to enthesis development *in vivo* but these cues have not been widely explored in engineered entheses and their individual contribution to maturation is largely unknown. Our objective here was to investigate how slow stretch, mimicking ACL growth rates, and intermittent cyclic loading, mimicking muscle activity, individually drive enthesis maturation in our system so to shed light on the cues governing enthesis development, while further developing our engineered replacements. Interestingly, we found these loads differentially drive organizational maturation, with slow stretch driving improvements in the interface/enthesis region, and cyclic load improving the ligament region. However, despite differentially affecting organization, both loads produced improvements to interface mechanics and zonal composition. This study provides new insight into how mechanical cues differentially affect enthesis development, while producing some of the most organized engineered enthesis to date.

## 1. Introduction

Ligaments and tendons connect to bone via structurally complex attachment sites known as entheses. Entheses are compliant fibrocartilaginous attachments comprised of gradients in organization, composition, mineralization, and cell phenotype. Organizationally, entheses are characterized by a transition from highly aligned collagen fibers in the ligament to more diffuse collagen fibers with oblique insertions into the bone [1–3]. These diffuse and oblique collagen insertions at the ligament-to-bone interface are hypothesized to be critical to reducing stress concentrations, while increasing toughness and compliance of the bone attachments [4–8]. In addition to gradients in collagen organization, gradients in matrix composition and tissue mineralization also contribute to proper translation of loads between the highly dissimilar elastic ligaments and stiff bone [9–11].

The anterior cruciate ligament (ACL) is the major stabilizer of the knee, connecting the femur to the tibia via entheses. In the U.S. alone, there are approximately 225,000 ACL injuries each year [12]. Following injury, enthesis healing is limited, and neither graft repair nor engineered replacements restore the organizational and compositional gradients necessary for proper long-term enthesis function, contributing to high re-tear rates [9–11]. In fact, of the 135,000 ACL reconstructions performed annually in the U.S. [12], approximately 10-25% retear within five years leading to further ACL revision surgery [13–16]. Additionally, subpopulations such as adolescents, young women, and individuals returning to soccer have been reported to be 1.5-3 times more likely to undergo ACL revision surgery within two years of repair [17,18]. Failure of ACL grafts has been shown to occur largely at the enthesis due to decreased bone mineral density and unorganized scar tissue formation at the insertion site [9,10]. Furthermore, ACL revision surgery has a retear rate of 2-25%, a clinical failure rate of approximately 20-30%, and is complicated by concomitant pathologies following ACL reconstruction such as cartilage damage, meniscal defects, tunnel-site bone deficiencies, or peripheral joint laxity [17–19]. Therefore, there is a need to develop engineered replacements with functional entheses or drive regeneration of entheses to improve ACL repair outcomes and avoid further costly, painful revision surgery and associated concomitant pathologies.

Previously, we developed a static tensile-compressive culture system (**Fig. 1A**) that guides neonatal bovine ACL fibroblasts in high-density collagen gels to form highly aligned collagen fibers in the mid-section and enthesis-like tissue on the ends. Specifically, compressive boundary restraints at the edge of the constructs produce a tensile-compressive environment, which guides cells to produce 3 unique zones of organization, composition, and cell phenotypes, recapitulating early postnatal-like entheses [12]. Constructs begin culture completely unorganized, and by 2 weeks develop aligned collagen fibers in the middle, perpendicular fibers in the transition, and unorganized collagen under the clamp, mirroring early postnatal-like enthesis development [12]. With more time in culture, organization, composition, and mechanical properties continue to mature; however, tissue maturation is largely confined to the early postnatal stage of enthesis development [2,20]. These constructs are some of the most organized engineered entheses to date and have promise as potential engineered replacements; however, driving organization beyond early postnatal development is critical for proper function *in vivo*.

**Figure 1:**
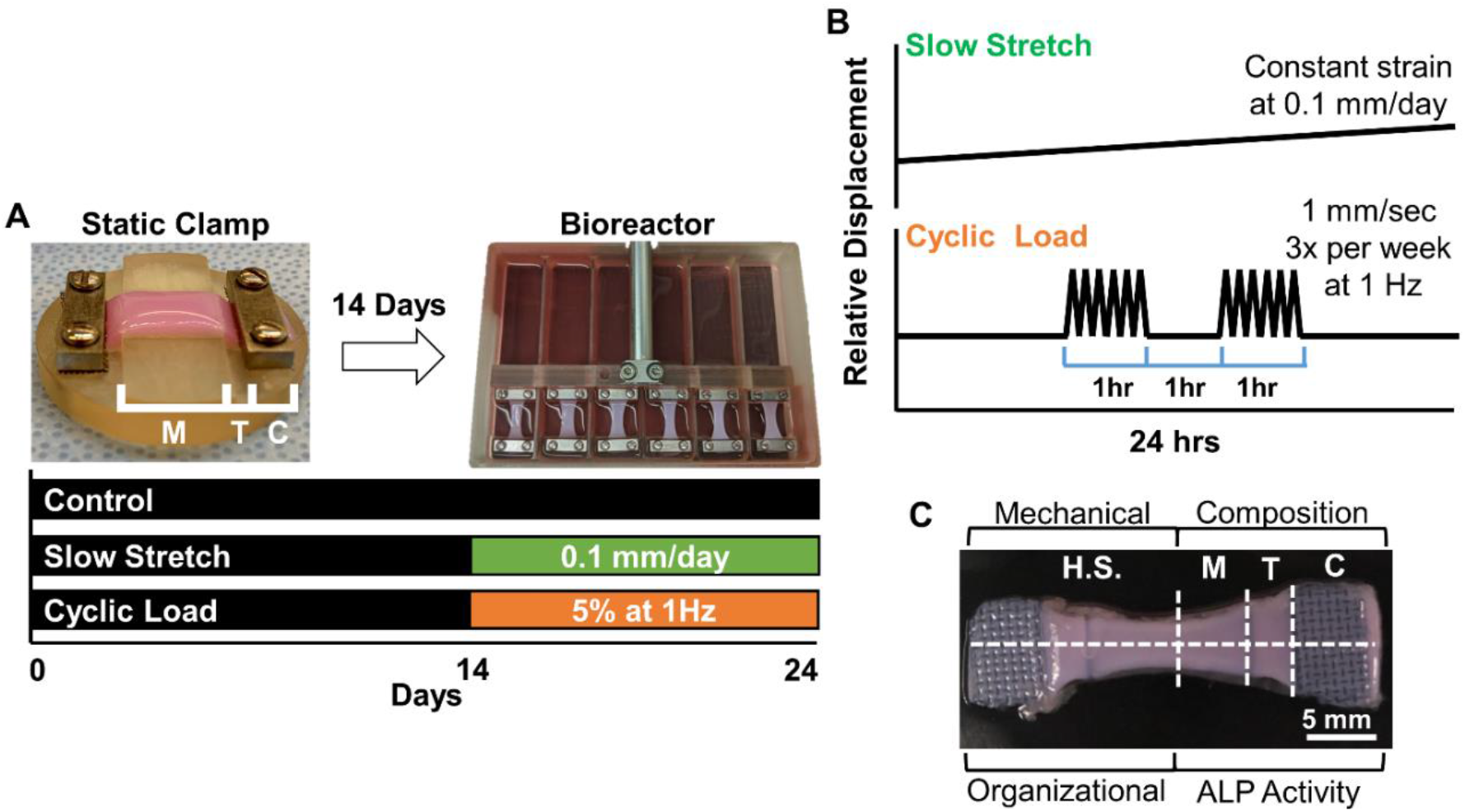
Depiction of culture conditions. (A) Cell-seeded high-density collagen gels were cultured in static clamping devices for 14 days to drive early postnatal-like enthesis development. After 14 days, loaded constructs were transferred to a modified CellScale tensile bioreactor and loaded for an additional 10 days, while control constructs remained statically clamped. (B) Depiction of loading regimes for slow stretch and cyclic load. Slow stretch was applied at 0.1 mm/day for 10 days, resulting in a final displacement of 1 mm. Cyclic load constructs were loaded with an established intermittent loading regime, 3 times a week (MWF), ensuring a total of 5 cycles in 10 days. Each cycle consisted of 1 hour on, 1 hour off, 1 hour on at 5% strain (1 mm) at 1 Hz. (C) Depiction of tissue sectioning performed upon removal from culture. Tissue was allocated as half-length sections (H.S.) for mechanical and organizational analysis, or into middle (M), transition (T), and clamped (C) sections for composition and alkaline phosphatase (ALP) activity analysis.

As the enthesis matures *in vivo*, collagen fibers shift from being largely perpendicular to the ligament, to being parallel with the ligament, with a more diffuse orientation and oblique insertions into the bone [1,2,20]. As discussed previously, this final collagen organization is hypothesized to be critical to reducing stress concentrations, while increasing toughness and compliance of the enthesis [4–8]. Dynamic mechanical cues such as ACL growth and cyclic muscle loading are necessary for enthesis maturation *in vivo* [21–23]. Early initiation of the enthesis has been shown to be independent of dynamic mechanical cues; however, enthesis maturation in later stages of development is driven largely by dynamic mechanical loading [22,24]. If these loads are not present, the enthesis fails to mature resulting in impaired collagen organization, fibrocartilage formation, mineralization, and tissue mechanics [21–25].

Most work assessing the role of mechanical loads on enthesis development has focused on the effects of cyclic muscle activity [21,24,26–28]. While these analyses are important, the developing enthesis experiences complex loading patterns with multiple mechanical cues, including slow growth elongation and compression. It is challenging to separate these mechanical cues *in vivo*; therefore, it is unclear which mechanical signals lead to the mature enthesis [23,29]. Understanding how each load respectively drives enthesis maturation could aid in identifying potential targets for regenerating entheses after injury, develop optimal rehabilitation protocols to improve regeneration after repair, and produce clinically relevant replacements [11,30–32].

Just as mechanical cues are critical for proper maturation of native entheses, mechanical conditioning is well established to enhance the development of various engineered musculoskeletal tissues. Much of this work has focused largely on tendon and ligament proper and has been largely understudied for enthesis engineering. Work in enthesis engineering has primarily focused on the application of cyclic loads mimicking muscle activity [33–35]; however, this research has focused on prefabricated scaffolds or decellularized native tissue with organization already set. The application of cyclic load to these tissues elicited a strong response in the tendon mid-substance but had little effect on the enthesis and bone regions, possibly due to the stiffer prefabricated scaffolds or decellularized bone shielding cells from mechanical signals [36]. Additionally, slow stretch has been investigated in tendon tissue engineering and has been reported to improve fibril diameter, packing, and mechanics [37]; however, the application of slow stretch has not been investigated in engineered entheses. Our culture system, which mirrors enthesis development provides a novel ability to decouple the effects of slow growth elongation and cyclic loading to shed light on their individual contributions to enthesis maturation. Understanding the individual effects of slow growth elongation and cyclic muscle loading will lead to improved engineered ligament replacements with functional ligament-to-bone entheses.

The objective of this study was to investigate how slow elongation, mimicking ACL growth rate, and cyclic loading, mimicking muscle activity, individually drive enthesis maturation in our culture system so to better understand how mechanical cues govern enthesis maturation, while further developing our engineered ligament replacements. We hypothesize that slow stretch drives maturation in fibril-level organization and cyclic loading drives maturation at the fiber and fascicle-level, with both slow stretch and cyclic loading yielding overall improvements in hierarchical collagen organization, tissue mechanics, and matrix composition, ultimately resembling late postnatal to mature entheses.

## 2. Methods and Materials

### 2.1 Construct Fabrication

Cell seeded high density collagen gels were fabricated as previously described [12,38–41]. Briefly, ligament fibroblasts were isolated from neonatal (1-3 days old) bovine cranial cruciate ligaments (CCLs), the bovine analog of human ACLs [12,38–42]. Three isolations were performed, with each isolation consisting of 3-4 donors combined into a single cell suspension. Cells were seeded at 2800 cells/cm^2^ and expanded in Dulbecco’s Modified Eagle Medium (DMEM) with 10% fetal bovine serum (FBS), 1% antibiotic/antimycotic (A/A), 50 μg/mL ascorbic acid, and 0.8 mM L-proline, changed every 3 days. Cells were expanded 1 passage and then seeded into constructs.

Constructs were composed of high-density collagen gels produced as previously described [12,38,43]. Briefly, type I collagen was extracted from Sprague Dawley rat tail tendons (BIOIVT) and reconstituted at 30 mg/mL in 0.1% acetic acid [12,38,44]. The collagen solution was mixed with NaOH and PBS to initiate gelation and ligament fibroblasts were immediately mixed into the collagen solution to ensure cells were seeded evenly throughout the gel. The final solution was injected between glass sheets 1.5 mm apart and gelled at 37°C for 1 hour yielding sheet gels at 20 mg/mL collagen and 5 million cells/mL [12,38,43]. Each sheet gel was made from a single expansion of cells using unique batches of collagen to account for batch-to-batch variation. Using a die, approximately 4-5 uniform constructs (30 × 8 mm) were cut from each sheet gel and allocated across timepoints and experimental groups.

### 2.2 Culture Conditions and Mechanical Loading

All constructs were clamped into static clamping devices on day 1 as previously described [12,38]. Steel mesh was placed between the clamp and the construct to facilitate nutrient diffusion under the clamp, as well as to prevent construct slippage throughout culture [12,38]. Constructs were cultured in the static clamping device for 14 days to induce early postnatal-like organization as previously reported [12]. After 14 days, mechanically loaded constructs were transferred to a modified CellScale tensile bioreactor and stimulated with either slow stretch or intermittent cyclic loading for 10 days (**Fig. 1A**). Slow stretch constructs were stretched at a rate of 0.1 mm/day, mimicking postnatal ACL growth rate (**Fig. 1B**) [45]. Cyclic load constructs were loaded with an established loading regime of 5% strain (1 mm) at 1 Hz for 1 hour on, 1 hour off, 1 hour on, 3 times per week (Monday, Wednesday, Friday, **Fig. 1B**), to mimic cyclic muscle activity [39]. Control constructs remained statically clamped throughout culture. Of note, both slow stretch and cyclic load groups underwent 1 mm of total applied strain throughout culture; however, this loading was applied at drastically different timescales of 10 days for slow stretch and 1 second for cyclic load. Constructs were cultured in the same media composition that cells were expanded in. Conditioned media changes were performed by removing and replenishing half of the media during each media change. These media changes were done directly before the application of load on Monday, Wednesday, and Friday throughout culture.

### 2.3 Post-Culture Harvesting and Analysis

At 2, 14, and 24 days constructs were removed from culture, weighed, and sectioned into half-length samples for organizational and mechanical characterization, or middle, transition, and clamped regions (M, T, and C) for analysis of zonal composition and ALP activity (**Fig. 1C**). The first timepoint was taken on day 2 to allow for 24 hours of clamped culture as a baseline of comparison [12,38]. Upon removal from culture, construct weights were measured and normalized to average construct weight at 2 days to assess change in mass throughout culture [12,38]. Additionally, constructs were photographed at each timepoint and surface area between the clamps was determined using FIJI (NIH) to track contraction (N = 6-12) [12]. Due to significant contraction, not all sections for analysis could be obtained from loaded constructs (**Fig. 1C**), resulting in a greater number of constructs produced for loaded groups (N = 12) compared to control static constructs (N = 6).

#### 2.3.1 Zonal Collagen Organization

Assessment of zonal hierarchical collagen organization at the fibril (<1 μm), fiber (1-100 μm), and fascicle (>100 μm) length-scales were performed via scanning electron microscopy (SEM), confocal reflectance, and picrosirius red staining as previously reported [12,38]. Briefly, fully intact half-length sections containing middle, transition, and clamped regions were fixed in 10% formalin and stored in 70% ethanol. Middle, transition, and clamped regions of engineered tissues were compared to 4-5 months old immature bovine CCL ligament, enthesis, and bone regions, respectively. A total of 6 constructs per timepoint and condition were analyzed via confocal reflectance. Following confocal analysis, a subset of 24 day control, slow stretch, and cyclic load constructs were processed for SEM (N = 3) and picrosirius red staining (N = 3).

Confocal reflectance analysis was performed with a Zeiss 710 inverted laser scanning microscope and LD C-Apochromat 40x/1.1 W Corr M27 objective as previously described [12,38–41]. Briefly, a 405 nm laser was used to visualize collagen at 400-465 nm by collecting backscattered light through a 29 μm pinhole, while simultaneously a 488 nm laser was used to visualize cells via autofluorescence at 509-571 nm. Half-length constructs were imaged intact and flat with 3-6 representative images taken per tissue region (total 9-18 images per construct). All images were analyzed using FIJI to assess collagen fiber alignment direction and dispersion as previously described [12]. Briefly, the maximum angle of fiber alignment and degree of fiber dispersion were determined for each image (n = 3-6 per region). Values from each image within a tissue region were then averaged to determine the average angle of alignment and dispersion for the middle, transition, and clamped regions of each construct (N = 6). The reported angle of alignment and dispersion are the average and standard deviation of the construct values (N = 6).

SEM analysis of 24 day constructs and native tissue was performed using a Hitachi SU-70 FE-SEM as previously described [12,43]. Prior to SEM imaging, the top layer of tissue was removed to better expose fibrils in the mid-substance of the construct using a cryostat. Briefly, constructs were washed twice in PBS for 5 mins, embedded flat in optimal cutting temperature (OCT) compound on a glass slide, frozen in a cryostat, and 10-12 slices (∼5 μm thick, at a 2.5° angle) were removed from the top layer of the constructs. Constructs were then thawed, washed twice in PBS, serially shifted from PBS to 100% ethanol, critical point dried, mounted flat with cryostat cut-side up on aluminum mounts with double sided carbon conductive tape, and coated in ∼5-6 nm of platinum. Constructs were imaged at 15 mm working distance, 5 kV, and 30,000x magnification, with 3-6 images taken per construct region. Half-length constructs were imaged and analyzed via FIJI to assess collagen fibril alignment direction and dispersion as described for confocal images.

Histological analysis was performed as previously described [12,38]. Briefly, fixed half-length 24 day samples and native bovine CCL-to-bone tissue were embedded into paraffin blocks, sectioned along the frontal plane, and stained with picrosirius red. Constructs were imaged using a Nikon Eclipse Ts2R inverted microscope and Nikon Plan Fluor 10x/0.30 OFN25 Ph1 DLL objective under linear polarized light at 10x to observe fascicle level collagen organization.

#### 2.3.2 Mechanical Analysis

Mechanical testing of the transition region was performed on an EnduraTEC 3200 System using a 250 g load cell as previously described [12,38,39,41]. Briefly, upon removal from culture, half-length sections (N = 6) were frozen at -20°C for storage. Samples were thawed prior to testing in PBS with EDTA-free protease inhibitor, measured, and placed into testing clamps. Constructs were secured into the testing clamps such that the full transition region was in between the two clamps. Constructs were loaded to failure at a rate of 0.75% strain/second, assuming quasi-static loading, and ensuring failure between the grips.

The resulting stress-strain data was analyzed using a custom least square linear regression-based MATLAB code to determine mechanical properties including stiffness, elastic modulus, ultimate tensile strength (UTS), strain at failure, toe region modulus, and the transition stress and strain (**Supplemental Fig. 1**). The linear elastic and toe region of the stress-strain curves were fit with linear regression and the resulting moduli were found as the maximum slope for each linear fit with an r^2^ > 0.999. Stiffness, calculated as elastic modulus times cross-sectional area and divided by gauge length, was also determined to account for differences in cross-sectional area between engineered constructs. The UTS and strain at failure were defined as the maximum point on the stress-strain curve. Transition stress and strain, defined as the transition point between the linear toe region and the linear elastic region, was defined as the closest point on the stress-strain curve to where the elastic and toe region linear fits intersected.

#### 2.3.3 Biochemical Analysis

Zonal analysis of DNA, GAG, collagen content, and alkaline phosphatase (ALP) activity were performed as previously described [12,38–41,46]. Briefly, constructs were sectioned into middle, transition, and clamped regions upon removal from culture. Tissue regions for DNA, GAG, and collagen analysis were weighed wet, frozen, lyophilized, weighed dry (DW), and digested in 1.25 mg/mL papain solution at 60°C for 16 hours (N = 6). DNA, GAG, and collagen content were then determined via Quantifluor dsDNA assay kit (Promega), modified 1,9-dimethylmethylene blue (DMMB) assay at pH 1.5 [47], and a modified hydroxyproline (hypro) assay [48]. Construct sections for ALP activity analysis were placed in 0.2% Triton X-100 immediately following post-culture sectioning and frozen at -80°C (N = 6). Samples were thawed, sonicated, and ALP activity was determined via a modified *p*-nitrophenyl phosphate assay [12,46]. Since constructs were made from high-density collagen gels seeded with ACL fibroblasts, DNA and collagen content were normalized to sample dry weight due to regional differences in wet weight caused by clamping. GAG and ALP are only added to the construct by cells and therefore are reported as normalized to DNA to account for differences in zonal cell distribution and activity.

#### 2.3.4 Immunofluorescence

Immunofluorescence was performed to assess localization of aggrecan and types I, II, and X collagen as previously described [12]. Briefly, day 24 engineered and demineralized bovine CCL-to-bone samples (N = 3) were fixed, embedded in paraffin blocks, and sectioned along the frontal plane. Sections were treated with proteinase K to retrieve antigens, blocked with 5% goat serum, and incubated overnight with primary polyclonal rabbit antibodies in 5% goat serum at 1:150 dilution for types I, II, and X collagen (Abcam AB34710, Abcam AB 34712, and GeneTex GTX37732, respectively), and aggrecan (GeneTex GTX54920). Negative controls were incubated overnight in 5% goat serum. All sections were then incubated with goat-anti-rabbit IgG secondary antibody labeled with Alexa Fluor 488 (Invitrogen A11008) at 1:200 dilution for 2 h and cross labeled with DAPI at 1:1000 dilution. Labeled sections were imaged on a Nikon Eclipse Ti2-E inverted microscope and Nikon Plan Fluor 10x/0.30 DIC L objective.

### 2.4 Analyzing Effects of Prolonged Cyclic Loading

Prolonged cyclic loading was investigated to assess if the additional loading time could drive further tissue maturation and account for differences observed at 24 days between slow stretch and cyclic load groups. Constructs were cultured as described in *2.2 Culture Conditions and Mechanical Loading*; however, constructs were treated with the cyclic loading regime for 20 days, compared to 10 days, for a total time in culture of 34 days. Following culture, all analyses described above were repeated for constructs receiving 20 days of cyclic load and compared to constructs receiving 10 days of cyclic load.

### 2.5 Statistics

Normality of the data and presence of outliers were assessed via Shapiro-Wilk tests and Q-Q plots in SPSS. Following confirmation of normality, data for fiber and fibril image analyses, mechanical analysis, and biochemical analysis were analyzed by 2 and 3-way ANOVA and Tukey’s *t*-test for *post hoc* analysis with *p* < 0.05 as significant. Data comparing middle, transition, and clamped regions of the same scaffolds were treated as repeated measures and therefore to assess differences between tissue regions repeated measures ANOVAs were used. All data are represented as mean ± standard deviation (S.D.).

## 3 Results

### 3.1 Gross Morphology Analysis

Gross analysis of constructs once removed from culture revealed that both slow stretch and cyclic load groups had similar changes in tissue morphology over 24 days in culture (**Fig. 2A**). Loading appeared to accelerate contraction, with slow stretch and cyclic load constructs having a significant decrease in percent area between the clamps to 40-45% of original area, compared to static controls contracting to ∼85% by 24 days (**Fig. 2B**). However, despite differences in percent area, all groups resulted in a similar significant decrease in weight by 24 days, reaching ∼25-30% of their original weight (**Fig. 2B**).

**Figure 2:**
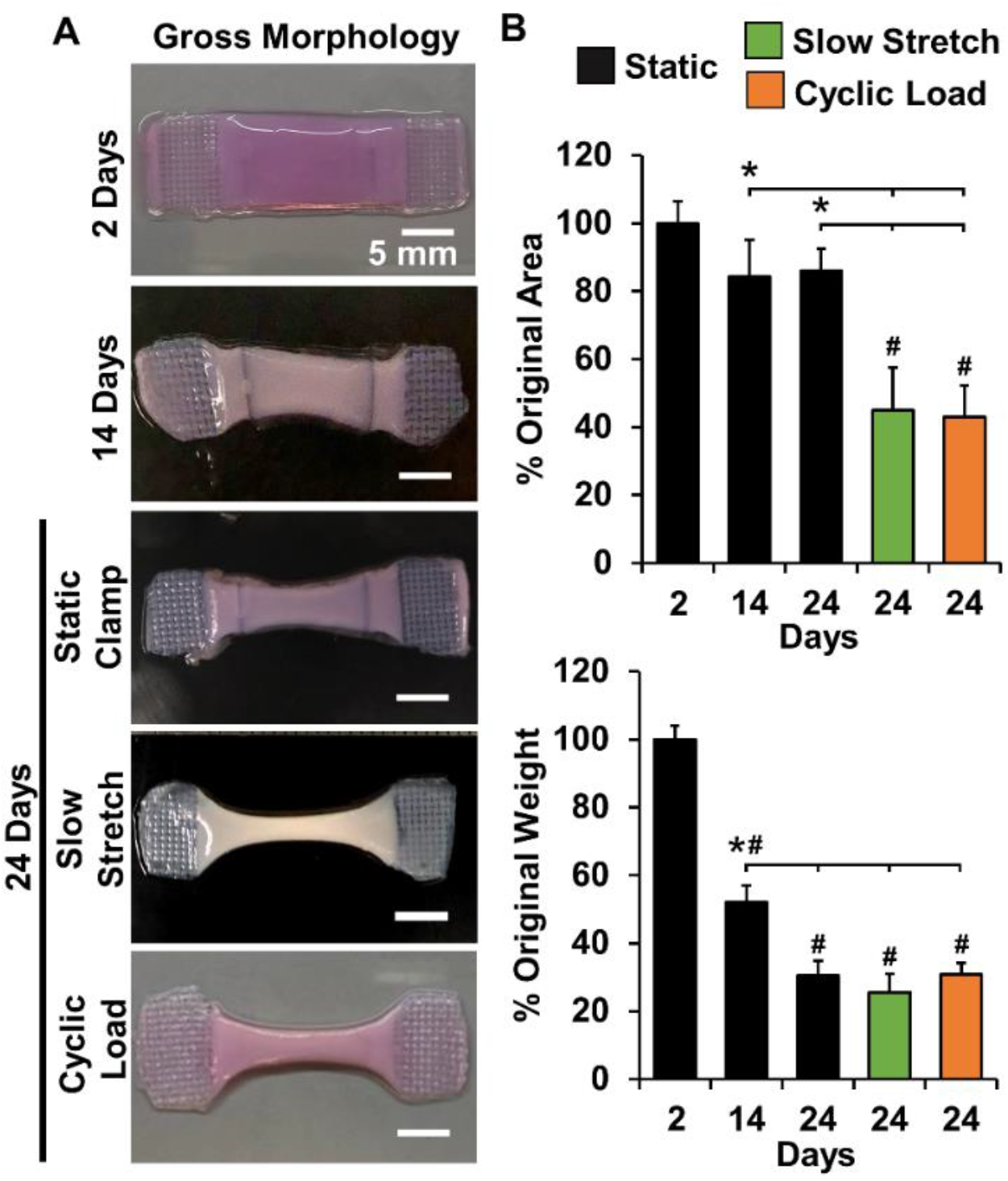
(A) Photographs of constructs at each timepoint depicting contraction throughout culture. (B) Construct percent area and weight throughout culture compared to constructs at 2 days. N = 6 static constructs, and N = 12 for loaded constructs. Significance compared to *bracket or ^#^day 2.

### 3.2 Hierarchical Collagen Organization Analysis

#### 3.2.1 Zonal Collagen Fiber Length-Scale Organization

Confocal reflectance revealed all groups development of 3 zones of unique collagen organization in the middle, transition, and clamped regions by 24 days in culture. Loading further increased collagen organiztion compared to static controls; however, the loading conditions differentially drove regional maturation (**Fig. 3A**). At the start of culture, constructs were composed of unorganized collagen across all three regions. By 14 days of static culture, constructs developed early postnatal-like enthesis organization with aligned parallel collagen fibers in the mid-section, perpendicular fibers in the transition (in relation to fibers in the middle region), and largely unorganized collagen under the clamp (**Fig. 3A**) [3,12,20]. By 24 days, static controls maintained this organization, while loaded constructs differentially drove further maturation. Specifically, slow stretch primarily drove maturation within the transition region, resulting in a shift from more perpendicular aligned collagen fibers to more direct, diffuse fiber insertions, similar to those seen in more mature native entheses (**Fig. 3A** green box). Interestingly, cyclic loading did not drive this shift in fiber orientation in the transition, however it did appear to drive further maturation in the mid-section, producing improved fiber and fascicle formation compared to static controls and slow stretch constructs (**Fig. 3A** orange brackets).

**Figure 3:**
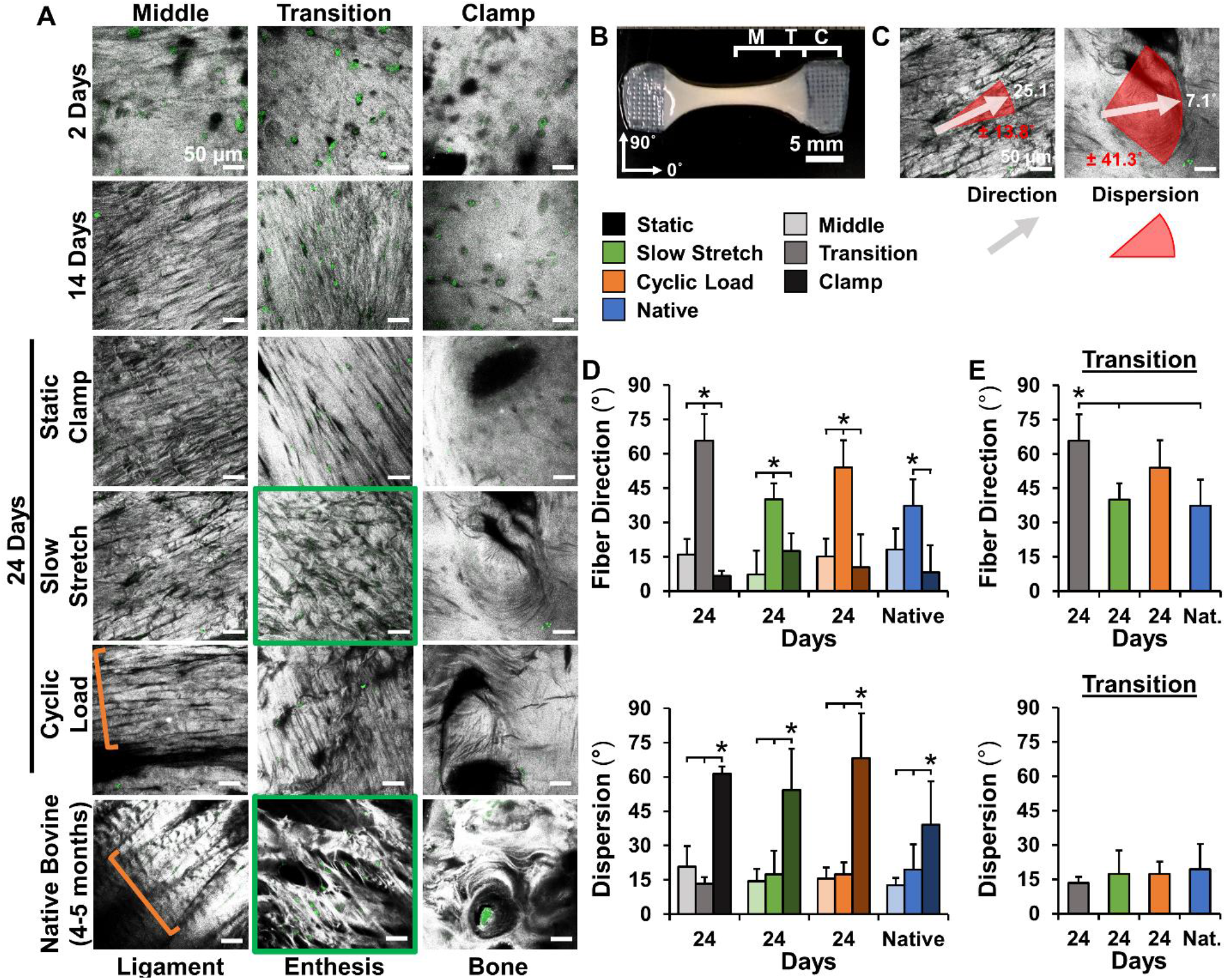
Slow stretch and cyclic load differentially drive zonal collagen fiber organization in engineered constructs. (A) Confocal reflectance reveals constructs develop aligned fibers in the middle, perpendicular fibers in the transition, and unorganized matrix under the clamp by 14 days. By 24 days, slow stretch improved maturation in the transition region (denoted by green box) and cyclic load improved maturation in the middle region (orange brackets depict fascicle formation). (B) Depictions of middle (M), transition (T), and clamped (C) tissue regions and angle orientation for image analysis. Confocal images were analyzed to determine angle of alignment and fiber dispersion in each region. (C) Example of alignment and dispersion analysis overlaid on confocal images of highly aligned images with low dispersion and unorganized regions with high dispersion. Image analysis of collagen fiber alignment and dispersion at 24 days compared to immature bovine entheses for D) all regions or E) just transition/insertion region. (E) Analysis of the transition region reveals that slow stretch resulted in a similar insertion angle to immature bovine ligament-to-bone insertions. 3-6 images were averaged per region for each construct or native tissue sample to determine construct averages. The reported angle of alignment and dispersion are the average and standard deviation of the construct averages for each tissue region (N=6 for engineered tissues and N=3 for native tissue). Significance compared to *bracket (p < 0.05).

Image analysis of collagen fiber organization at 24 days confirmed that slow stretch drove a shift in collagen fiber organization in the transition region compared to static controls and cyclic load constructs. By 24 days, all groups developed highly aligned collagen fibers with low dispersion in the middle and transition (aligned at ∼5-20° in the mid-section and 40-70° in the transition), and unorganized collagen with large dispersion of 55-70° under the clamp, similar to the regional organization observed in immature bovine CCL-to-bone insertions (**Fig. 3B-D** and **Supplemental Fig. 2A**). However, when looking specifically at the transition region, slow stretch constructs had a significant reduction in alignment direction to 40° ± 12°, similar to native immature entheses with a measured fiber alignment of 37.3° ± 11.6° in the transition (**Fig. 3E**). Comparatively, fiber alignments in the transition region for static controls and cyclic load constructs were 65.8° ± 7.1° and 54.0° ± 11.4°, respectively.

#### 3.2.2 Zonal Collagen Fibril Length-Scale Organization

Similar to fiber level organization evaluated by confocal microscopy, SEM revealed collagen organization at the fibril level was conserved for all groups at 24 days (**Fig 4A**). Fibril organization in the middle and clamped regions were largely similar amongst all groups with highly aligned fibrils in the middle region and unorganized fibrils under the clamp, resembling organization in immature bovine ligament and bone regions. However, slow stretch again drove a shift in the transition region from the more perpendicular fibril organization observed in static control and cyclic load constructs to more direct fibril insertions, mirroring fibril organization observed in immature bovine entheses (**Fig 4A**).

**Figure 4:**
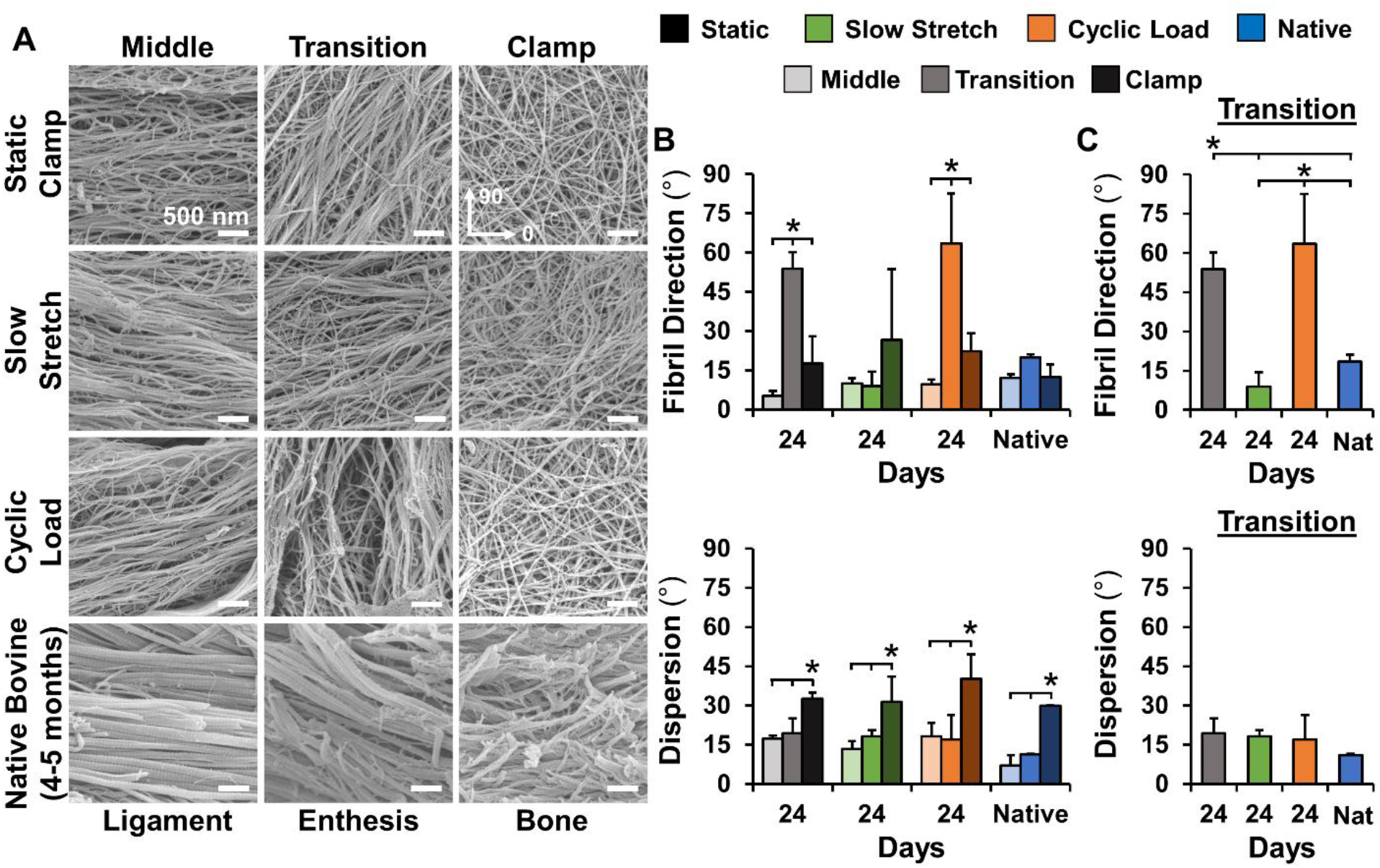
Slow stretch or cyclic load differentially drive zonal fibril level organization. (A) SEM imaging reveals that fiber level organization was conserved at the fibril level in all tissue regions at 24 days with slow stretch producing more direct fibril insertions in the transition. Image analysis of collagen fibril alignment and dispersion at 24 days compared to immature bovine tissue for B) all regions or C) just transition/enthesis region. (C) Analysis of the transition region again reveals that slow stretch produced a similar insertion angle as immature bovine ligament-to-bone enthesis. 3-6 images were averaged per region to determine construct averages (N = 3). The reported angle of alignment and dispersion are the average and standard deviation of the construct averages for each tissue region. Significance compared to *bracket (p < 0.05).

Image analysis of fibril level organization at 24 days confirmed that slow stretch drove a significant shift in fibril organization in the transition region compared to static controls and cyclic load constructs (**Fig. 4B**). By 24 days, all groups developed highly aligned collagen fibrils in the mid-section (aligned at ∼5-10°), and more unorganized collagen fibrils in the clamped region (dispersion of ∼30-40°). When assessing the transition region specifically, static control and cyclic load constructs had fibril alignments of 53.9° ± 6.2° and 63.5° ± 19.1°, respectively, while slow stretch constructs had a fibril alignment angle of 8.9° ± 5.6° which more closely mirrored the fibril organization measured in immature bovine entheses of 19.9° ± 1.1°.

#### 3.2.3 Zonal Collagen Fascicle Length-Scale Organization

Polarized light microscopy of picrosirius red stained sections at 24 days revealed that hierarchical collagen organization observed at the fibril and fiber level was also observed at the fascicle length-scale for all groups (**Fig. 5**). Within the middle region, both slow stretch and cyclic loading drove development of larger collagen fibers and early fascicle organization, compared to static clamped constructs. However, similar to the fiber level, cyclic loading drove further maturation in the middle region, developing larger fascicles with increased crimp formation. In the transition region, organization largely mirrored fiber and fibril level organization with slow stretch constructs having more direct, parallel insertions, similar to immature bovine transitions, while static controls and cyclic load constructs maintained more perpendicular fibers. Under the clamp, all groups developed collagen aligned circumferentially around pores created by the mesh, resembling collagen oriented around pores in native bone.

**Figure 5:**
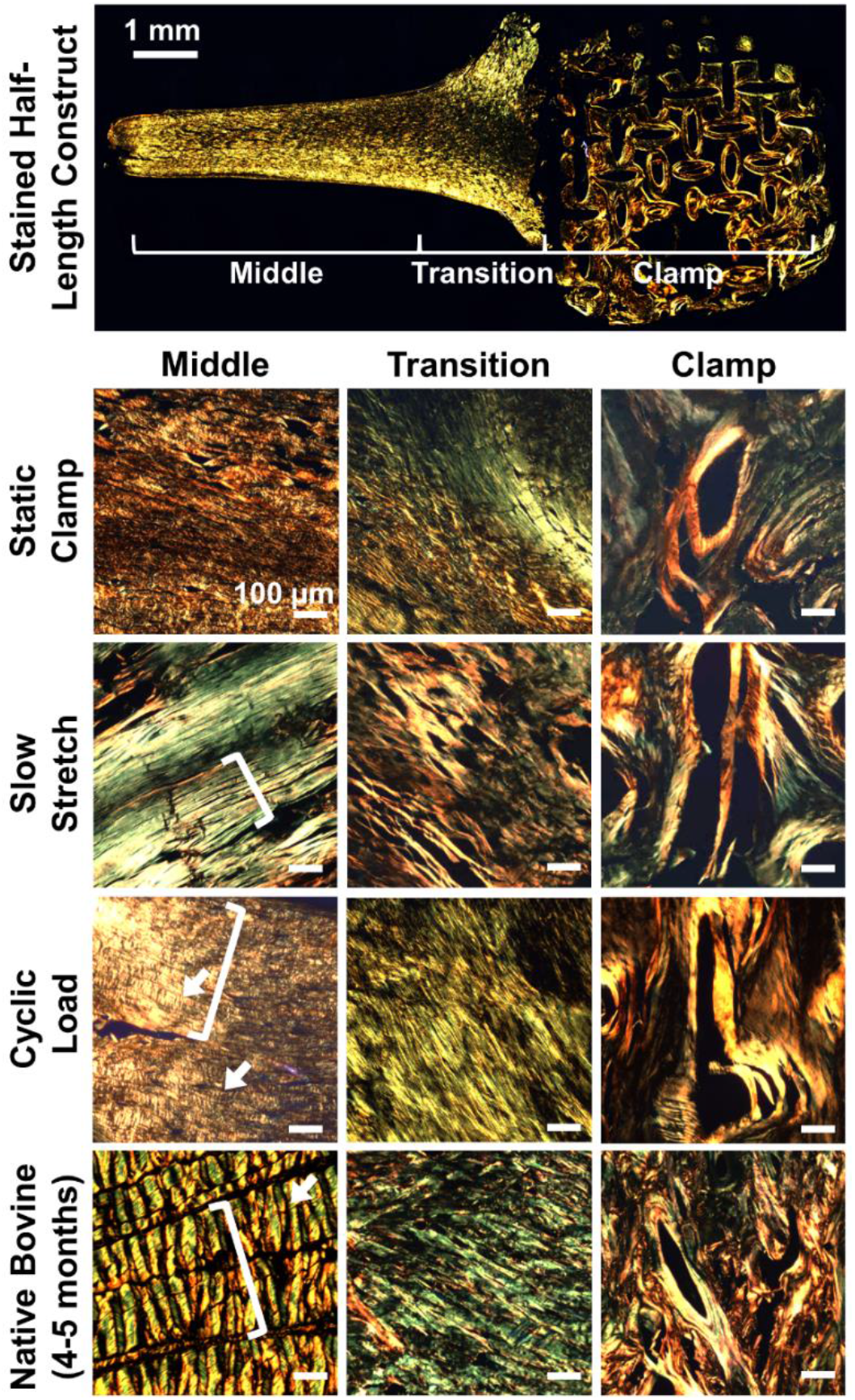
Organization at the fascicle length-scale evaluated by picrosirius red staining imaged with polarized light. Slow stretch produced more direct fiber insertions in the transition, similar to immature bovine enthesis, and cyclic load results in larger fascicle-like organization with early crimp formation, similar to immature bovine ligament. Brackets denote fascicle formation; arrows denote crimp.

Collectively, analysis of the fibril, fiber, and fascicle length-scales demonstrated the development of hierarchical, zonally organized collagen. Most notably, at all levels of organization, slow stretch drove a shift in the transition region from more perpendicular collagen alignment to more direct fiber insertions. This shift mirrored the observed organization of immature bovine entheses, indicating further tissue maturation compared to static control and cyclic load constructs. Additionally, at the fiber and fascicle length-scales cyclic loading appeared to drive improved fiber and early fascicle formation in the middle region, mirroring the organization of immature bovine ligament.

### 3.3 Transition Region Mechanical Properties

Despite driving differential maturation in collagen organization, both slow stretch and cyclic load constructs yielded similar significant improvements in transition region tensile mechanics compared to static controls (**Fig. 6**). Specifically, both slow stretch and cyclic load constructs had a 2-3 fold increase over static controls in stiffness and elastic modulus, reaching ∼0.4 N/mm and ∼3.5-4.5 MPa, respectively (**Fig. 6B**). Slow stretch alone produced a significant increase in UTS compared to static controls; however, the increase in UTS for slow stretch constructs was not significantly different than the UTS for cyclic load constructs (**Fig.6C**). Interestingly, while static controls had a significant decrease in strain at failure by 24 days, slow stretch and cyclic load constructs did not have a decrease in failure strain, indicating both loaded constructs may yield a more compliant transition region compared to that of static controls.

**Figure 6:**
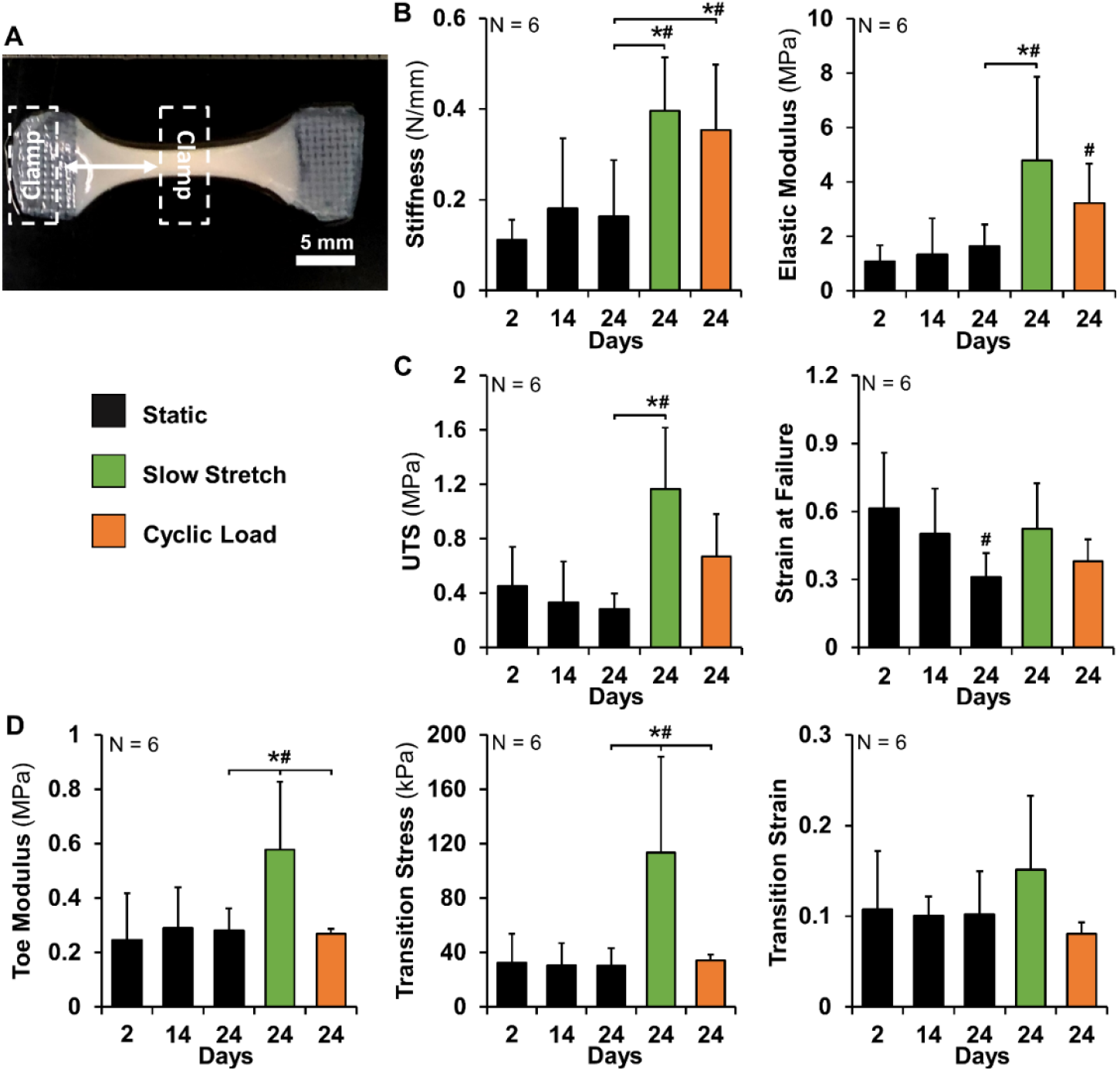
Slow stretch and cyclic load significantly improved tensile properties (A) across the transition. Slow stretch and cyclic load resulted in similar significant increases in B) stiffness and elastic modulus, and (C) similar increases in ultimate tensile strength (UTS), while remaining more compliant with increased strain at failure at 24 days. Slow stretch alone drove significant improvements to (D) toe region mechanics. Significance compared to *bracket or ^#^day 2.

Interestingly, only slow stretch constructs had significant differences in the toe region mechanics across the transition. By 24 days, slow stretch constructs had a 2 fold increase in the toe modulus over static controls and cyclic load constructs, reaching a modulus of 0.6 ± 0.3 MPa. (**Fig. 6D**). Finally, looking at the transition point where the stress-strain curve transitions from the toe region to the elastic region, slow stretch constructs had a 4-fold increase in the transition stress compared to static control and cyclic load constructs and no significant change in transition strain. Collectively, the toe region data indicates that slow stretch produces constructs with improved toe region toughness compared to static controls and cyclic loading.

### 3.4 Zonal Composition

Zonal collagen, DNA, and GAG concentrations were similar for all groups at 24 days (**Fig. 7A-C**). Specifically, collagen content, represented by hydroxyproline and normalized to dry weight, remained relatively constant throughout culture in all groups and tissue regions (**Fig. 7A**). DNA (normalized to dry weight) increased in the middle regions by 14 days and static controls maintained these levels through 24 days. The addition of load largely did not change the zonal concentrations of DNA, however both slow stretch and cyclic load did have an increase in DNA in the transition region, resulting in a significant increase compared to day 2, but not compared to day 24 static controls. Additionally, under the clamp all groups had no change in DNA throughout culture (**Fig. 7B**), however all groups did have a significant 2-3 fold increase in GAG/DNA by 24 days under the clamp compared to middle and transitions regions (**Fig. 7C**). Interestingly, ALP activity changed throughout culture and resulted in differences between tissue regions and groups at 24 days (**Fig. 7D**). All tissue regions had a similar significant increase in ALP activity by 14 days; however, in static controls, ALP activity levels returned to baseline in all regions by 24 days. The addition of 10 days of slow stretch produced a zonal distribution of ALP activity by 24 days with a significant increase in the transition region and under the clamp compared to static controls, as would be expected with further tissue maturation. Surprisingly, cyclic loading had the opposite effect and led to a zonal distribution with significant increases in ALP activity in the middle and transition regions while ALP activity returned to baseline under the clamp.

**Figure 7:**
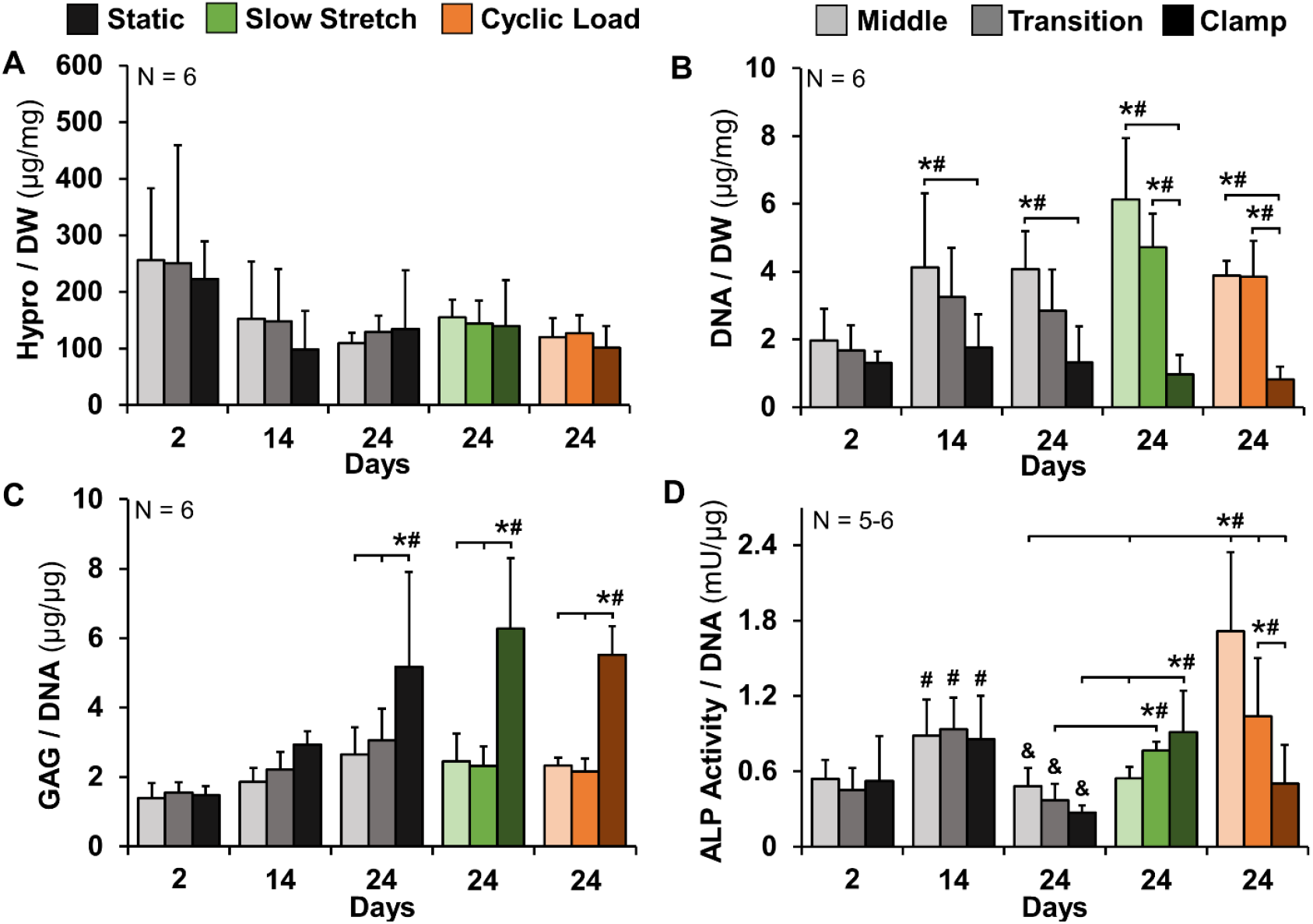
Loading had little to no effect on zonal collagen, DNA, and GAG, but drove differential zonal ALP activity. Zonal (A) Collagen content, represented by hydroxyproline (hypro), and (B) DNA normalized to dry weight (DW), (C) glycosaminoglycans (GAG) and (D) alkaline phosphatase (ALP) activity measured as nmol p-nitrophenol substrate catalyzed per minute (mU) normalized to DNA. Significance compared to *bracket or ^#^day 2 tissue region.

Constructs at 24 days were evaluated for spatial distribution of types I, II, and X collagen, and aggrecan (**Fig. 8**, negatives shown in **Supplemental Fig. 3**). As expected, type I collagen was dispersed uniformly throughout each tissue region for all groups (**Fig. 8A**). For all constructs, types II and X collagen were localized to the transition and clamped regions, similar to immature bovine enthesis and bone regions, with slow stretch and cyclic load yielding increased accumulation (**Fig. 8B-C**). Interestingly, both slow stretch and cyclic load appeared to drive increased type II and X collagen accumulation under the clamp; however, in the transition region slow stretch resulted in increased type II collagen, while cyclic load resulted in increased type X collagen. Aggrecan was throughout each tissue region for all groups; however, both load groups appeared to have increased aggrecan accumulation in the transition region compared to static controls, more closely resembling the aggrecan distribution observed in immature bovine entheses (**Fig. 8D**).

**Figure 8:**
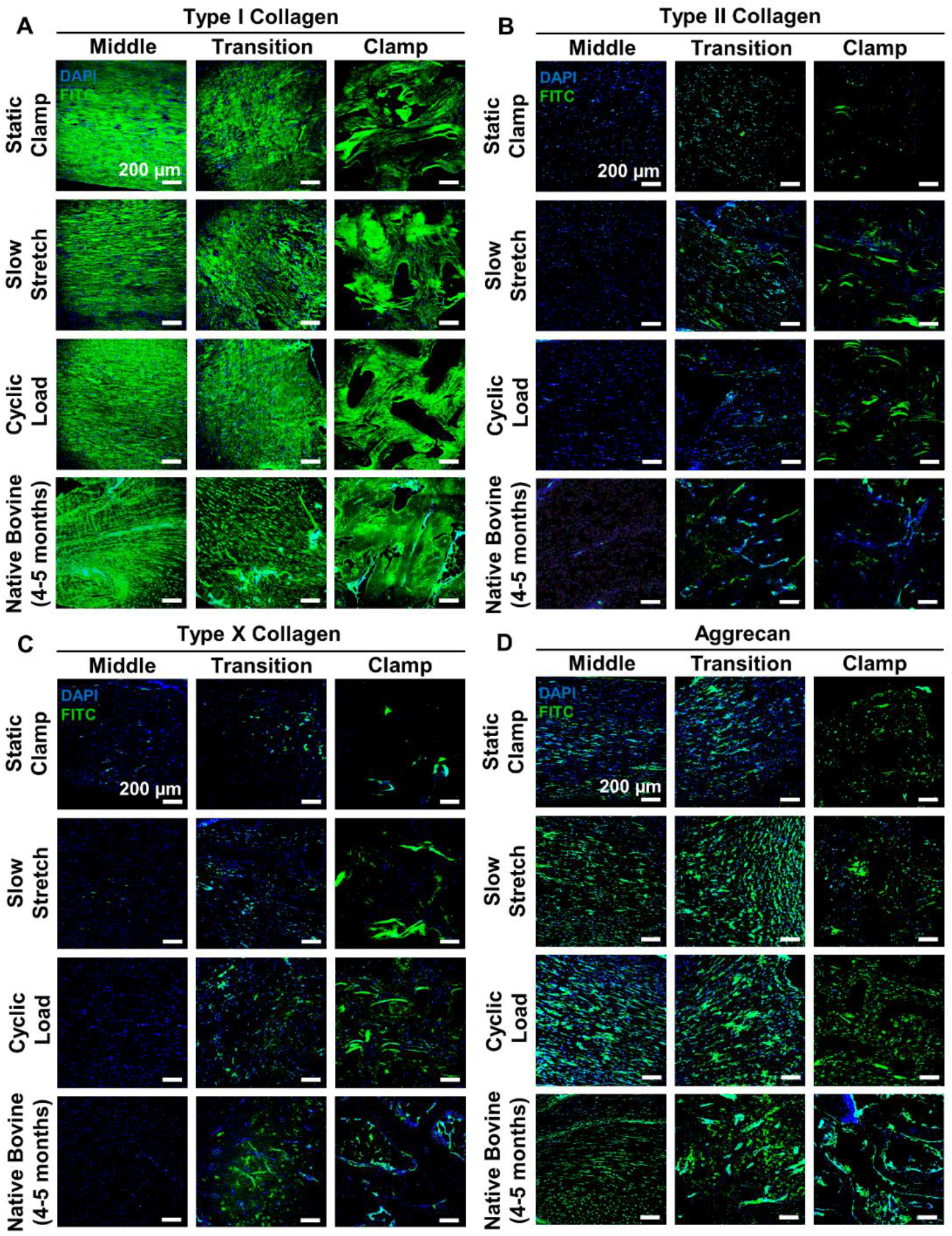
Loading increases zonal accumulation of type II collagen, type X collagen, and aggrecan. Immunofluorescence imaging of constructs at 24 days compared to late postnatal bovine ligament-to-bone tissue. Zonal accumulation of (A) Type I collagen; (B) Type II collagen; (C) Type X collagen; and (D) Aggrecan. FITC = respective protein, DAPI = nuclei, scale bars = 200 μm.

### 3.5 Effects of Prolonged Cyclic Load on Tissue Maturation

Interestingly, we found that slow stretch alone drove a change in collagen organization in the transition region resulting in a shift to more direct insertions. To investigate whether a longer culture period of cyclic loading could similarly drive this shift in organization, the experiment was repeated, and constructs were cyclically loaded for 20 days (34 day cyclic) and compared to previous 24 day samples (**Fig. 9A**). Gross morphology analysis revealed that 34 day cyclic constructs contracted more than 24 day cyclic constructs, with decreases in construct percent area and weight (**Fig. 9B**). Despite differences in contraction, zonal collagen organization at the fibril, fiber, and fascicle length-scale remained similar to 24 day cyclic constructs (**Fig 9C-D**). Image analysis of fibril alignment direction in the transition region revealed a shift in alignment from 63.5° at 24 days to 51.4° at 34 days (**Fig 9E**), however this shift was not significant, nor similar to that previously measured in slow stretch constructs and immature native tissue (fibril alignment ∼8-20° in the transition, **Fig 4C**). Further, there were no changes in alignment or degree of dispersion at the fiber length-scale with prolonged loading as well (**Fig 9F**). Collectively, this indicates that prolonged cyclic loading does not significantly drive further improvements in organization. Mirroring the relative lack of further maturation in 34 day cyclic constructs, tissue mechanical properties across the transition remained largely unchanged compared to 24 day cyclic load constructs (**Fig. 9G)**. However, although not significant, there appeared to be an overall decline in mechanics for 34 day constructs indicating that perhaps the additional load has a negative effect on the integrity of the transition region.

**Figure 9:**
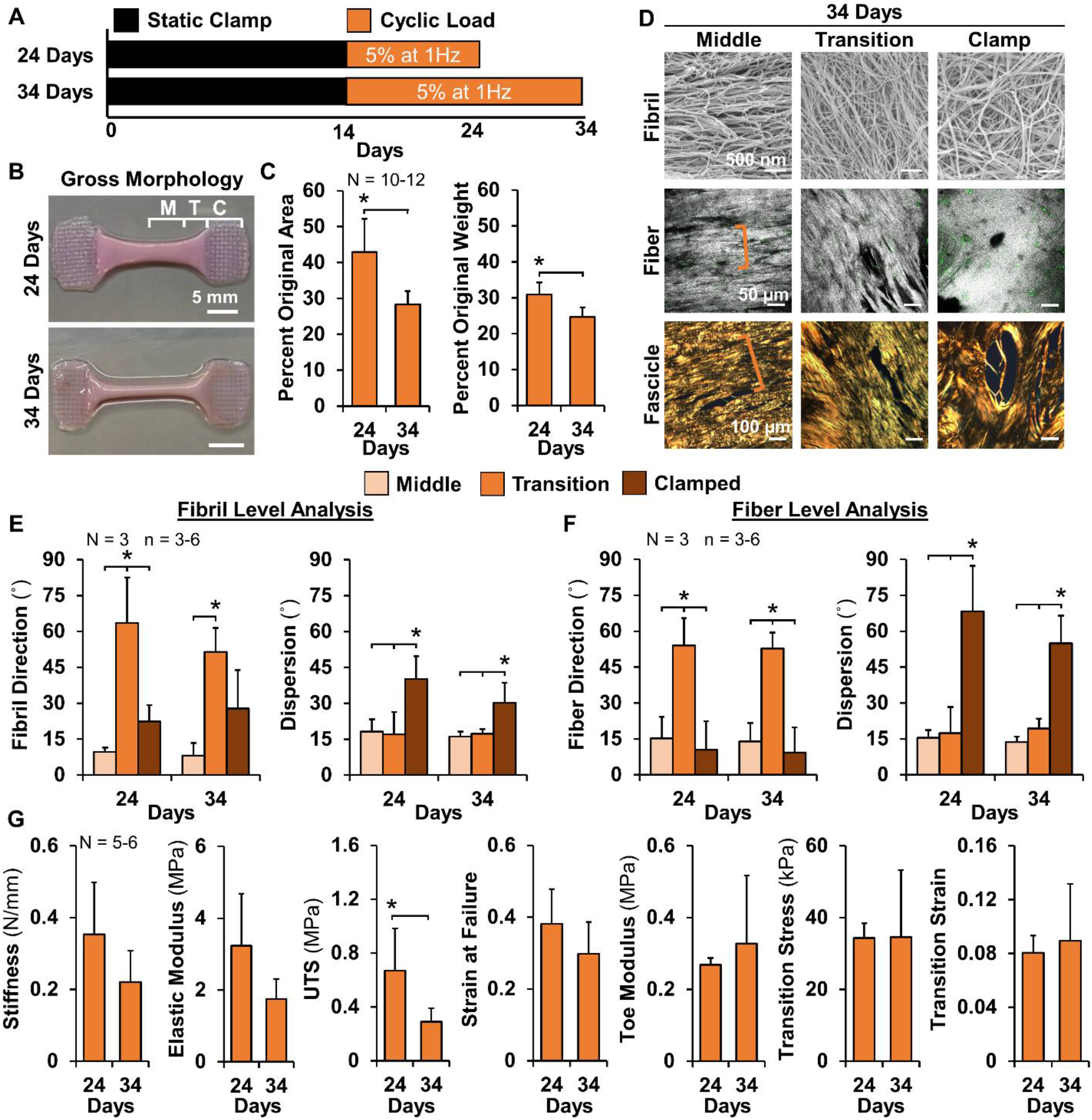
Prolonged cyclic loading does not alter organization or tissue mechanics. (A) 24 day cyclic load constructs were compared to constructs receiving 10 additional days of cyclic loading (34 day). (B) Gross morphology demonstrated that constructs were more tube-like in morphology at 34 days with a (C) significant decrease in construct percent area and weight. (D) There were no differences in hierarchical collagen organization at the fibril, fiber, and fascicle scales in 34 day cyclic constructs, analyzed via SEM, confocal reflectance, and picrosirius red respectively. (E) Fibril and (F) Fiber level image analysis confirmed that there were no changes in alignment or dispersion with 10 additional days of load. (G) 34 day cyclic load constructs also had no improvements in tensile mechanics across the transition region. Significance compared to *bracket (p < 0.05).

Similarly, there were minimal differences in zonal composition with prolonged cyclic loading. Specifically, there was no significant differences observed between 24 and 34 days for zonal collagen, DNA, or GAG accumulation (**Fig. 10A**). Interestingly, there was a significant decrease in ALP activity back to baseline levels in the middle and transition regions with prolonged cyclic loading, as well as the formation of a zonal distribution with significantly higher ALP activity observed under the clamp than in the middle and transition regions (**Fig. 10A**). This observation for cyclic load constructs at 34 days is similar to ALP activity observed in slow stretch constructs at 24 days. Additionally, there was little difference observed in types I, II, and X collagen, and aggrecan localization at 34 days compared to 24 day cyclic load constructs (**Fig. 10B**). Taken together, the data for 34 day cyclic loading suggests the need for combined or ramped loading to further drive maturation at the insertion site in tissue engineered constructs.

**Figure 10:**
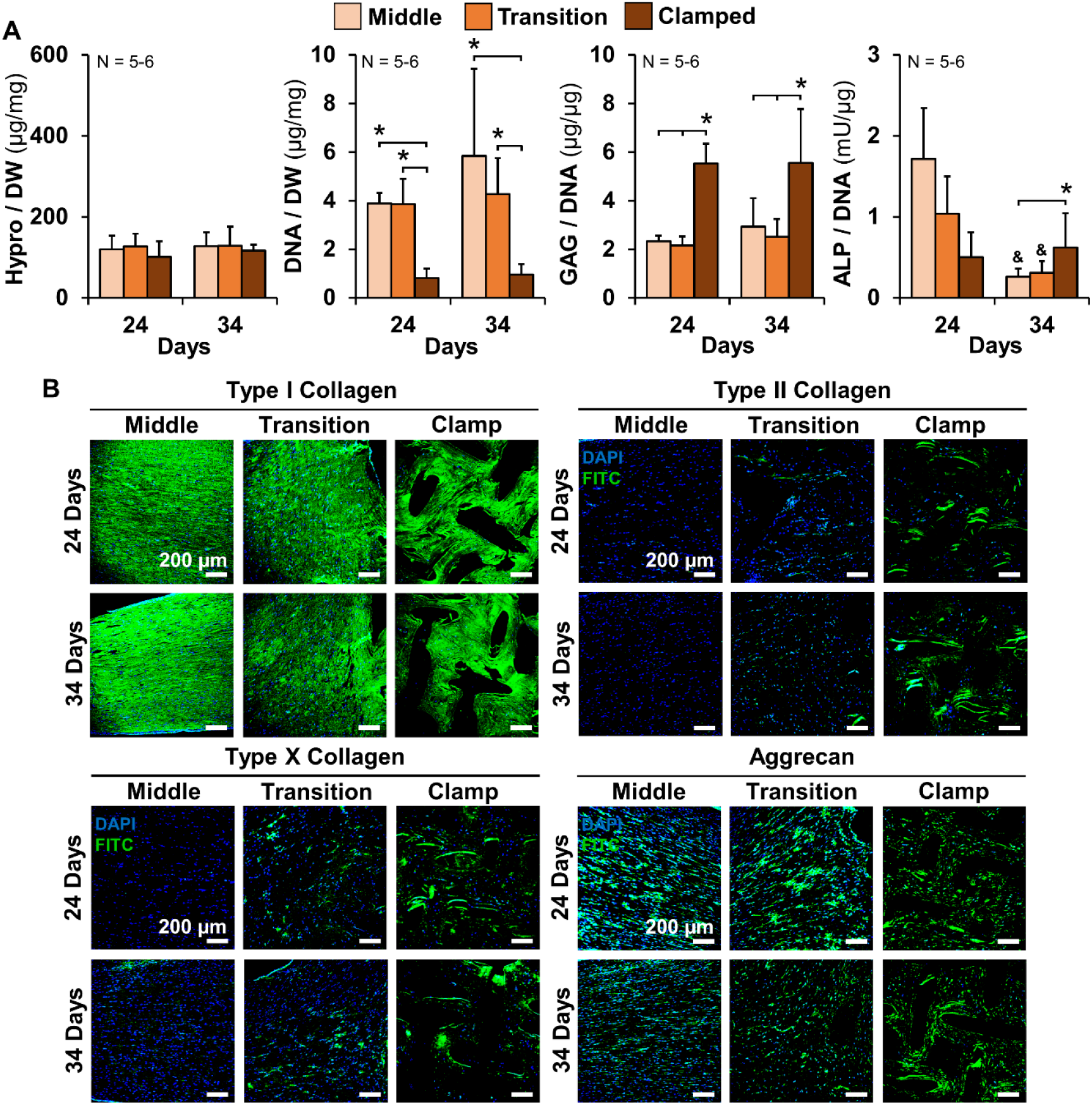
Prolonged cyclic loading has little effect on composition gradients. (A) Zonal Hydroxyproline (hypro) and DNA normalized to dry weight (DW), and glycosaminoglycans (GAG) and ALP activity measured as nmol p-nitrophenol substrate catalyzed per minute (mU) normalized to DNA. (B) Immunofluorescence imaging of type I collagen, type II collagen, type X collagen, and aggrecan, revealed no changes in protein accumulation or localization with prolonged cyclic loading. FITC = representative protein, DAPI = nuclei, scale bars = 200 μm. Significance compared to *bracket or ^&^24 day cyclic tissue region (p < 0.05).

## 4. Discussion

Fibrocartilaginous entheses are structurally complex tissues that attach elastic ligaments to stiff bone [9,10]. Entheses facilitate the load transfer between these highly dissimilar tissues via gradients in collagen organization, matrix composition, and mineralization [6–11]. Currently, these gradients necessary for the integrity of the entheses are not restored in ligament-to-bone healing, graft repair, or in engineered replacements. Therefore, there is a need to regenerate these gradients in engineered replacements or *in vivo* following injury and repair to create functional replacements and improve surgical outcomes. Recapitulating development is a promising way to produce functional entheses. Enthesis development is characterized by a shift in collagen organization in the fibrocartilage interface which is ultimately thought to give rise to the necessary mechanical function of the enthesis [2–8,20]. Initially, collagen organization at the interface is largely unorganized in the neonatal stage of development and shifts during early postnatal development to be oriented perpendicular to the ligament [3,20]. In late postnatal development, collagen organization shifts to being parallel with the ligament with more diffuse, oblique fiber insertions into the bone [1–3,20]. Maturation in this later stage of development has been shown to be driven largely by dynamic mechanical loading [21–24]; however, most work assessing these mechanical loads has focused on the effects of cyclic muscle activity [21,24,26–28]. There has been little consideration for slow growth elongation, partially due to the difficulty of decoupling these cues *in vivo* [23,29]. In this study, we aimed to investigate these cues by mimicking ACL growth rate and cyclic muscle activity in an *in vitro* culture system that mirrors native enthesis development [12]. This study demonstrates that slow stretch elongation and intermittent cyclic loading differentially drive zonal maturation in tissue engineered ligament-to-bone constructs, resulting in improved hierarchical collagen organization, tissue mechanics, and matrix composition, similar to late postnatal ACL-to-bone attachments.

Specifically, we found 10 days of slow stretch improved maturation in the transition region shifting hierarchical collagen fibrils and fibers to more direct insertion angles, closely mirroring the organization observed at the insertion site of late postnatal (4-5 months) bovine entheses (**Figs. 3-5**). Interestingly, only slow stretch drove this shift, whereas cyclic load constructs maintained more perpendicular oriented collagen in the transition, similar to static controls and early postnatal-like entheses [2,20]. However, contrary to slow stretch, 10 days of intermittent cyclic loading yielded improved maturation in the middle region, resulting in larger fibers and improved fascicle formation, similar to late postnatal bovine ligament (**Figs. 4** and **5**). Both slow stretch and cyclic load regimes stretched constructs the same amount (1 mm total); but at drastically different rates (0.1 mm/day vs 1 mm/sec). Thus, the change in zonal organization is not because of the total distance these constructs were stretched, but more so how cells sense and respond to different rates of load. Further, prolonged cyclic loading was still not capable of driving a similar shift in collagen organization at the interface compared to slow stretch (**Fig. 10**), indicating that slow growth elongation may be responsible for the shift of collagen from a perpendicular orientation to more direct, oblique insertions into the bone as entheses mature from early-postnatal to late-postnatal stages of development.

The shift in ligament-to-bone entheses organization observed between early- and late-postnatal development has largely been attributed to the effects of cyclic muscle activity [21,24,26–28]; however, these studies largely rely on techniques such as muscle paralysis to investigate the effects of cyclic muscle activity on the developing enthesis [49]. A limitation of this method is that the limb suffers arrested development and tissue atrophy as a result of muscle paralysis or removal [50], thereby also disrupting slow developmental strains associated with growth rates. Though it is difficult to deconvolute the effects of these mechanical cues *in vivo*, the results of this study suggest that the arrested growth, due to the loss of muscle activity, may be more responsible for the lack of organizational maturation at the insertion sites of developing entheses in previous studies *in vivo*.

Interestingly, despite slow stretch driving organizational improvements in the transition and cyclic loading driving improvements in the middle region, both loads produced similar 2 to 3-fold improvements in stiffness, elastic modulus, and UTS of the transition region (**Fig. 6A-B**). The moduli of slow stretch and cyclic load constructs surpassed the elastic modulus of 1 week old bovine entheses and cranial cruciate ligaments (analogous to human ACL, enthesis = 1 MPa, ligament = 1-3 MPa, [9,51]) and matched the UTS of the cranial cruciate ligament (0.8-2 MPa [51]). In addition to developing increased mechanics, both slow stretch and cyclic loading yielded constructs with higher failure strains compared to static controls. This data indicates that both slow stretch and cyclic load constructs developed increased strength over longer strains resulting in increased compliance and toughness of the transition region. This is characteristic of more mature entheses and has been shown to contribute to proper load transfer between ligament and bone [6,8,22].

Mirroring organizational changes, slow stretch alone drove increases in toe region mechanics across the transition region (**Fig. 6C**). Specifically, toe modulus and transition stress of slow stretch constructs were 3 to 4-fold higher than cyclic load and static constructs, while all constructs maintained a similar transition strain. In tendons and ligaments, the toe region is attributed to collagen fiber and fibril uncrimping or to the realignment of collagen fibers and fibrils during loading [4,5,30,52,53]; whereas in the enthesis, the toe region is attributed to fiber recruitment and the interactions of the fibers with the bone-ridge of enthesis attachments [54]. In slow stretch constructs, fibrils and fibers in the transition region are already oriented in the direction of the applied uniaxial tensile strain at the start of mechanical testing thus allowing for earlier recruitment and increased toe region stress. These organizational shifts likely account for the increased transition stress and toe modulus in slow stretch constructs, however, the lack of observed decrease in transition strain may implicate factors such as collagen fibril and fiber uncrimping, sliding, and shearing [30]. More in-depth mechanical and structural studies should be done to further investigate.

In addition to differentially regulating zonal organization, loading also differentially drove accumulation of key fibrocartilage matrix proteins in the transition and clamped regions. Specifically, loading had little to no effect on general DNA, GAG, and collagen concentrations; but it did differentially regulate accumulation of specific types of GAG and collagen, as well as zonal ALP activity. All constructs had a significant increase in GAG accumulation under the clamp (**Fig. 7C**), suggesting that clamping alone is sufficient to drive a shift toward a more cartilage-like tissue [55]. However, both slow stretch and cyclic load construct had increased aggrecan accumulation in the transition region compared to static controls (**Fig. 8C**), suggesting a shift toward more fibrocartilage-like tissue in the transition. Further, both slow stretch and cyclic load constructs had increased accumulation of type II and type X collagen in the clamped region and differential accumulation in the transition, with slow stretch resulting in increased type II collagen and cyclic load producing increased type X collagen. This suggests that slow stretch may contribute to the development of the fibrocartilage region of the enthesis and cyclic loading may contribute more to the mineralized fibrocartilage region, characterized by increased type II collagen and type X collagen, respectively [9,10].

Further, slow stretch drove a gradient in ALP activity with increased ALP activity under the clamp compared to middle and transition regions by 24 days (**Figure 7D**). ALP is a known marker in bone development and has been implicated in mineralization of the enthesis in later stages of development [2]. Interestingly, cyclic loading had the opposite effect at 24 days with increased ALP activity in the middle and transition regions; however, similar cyclic loading regimes have also previously been shown to acutely increase ALP activity in fibroblasts through 7 days of culture [56,57]. To our knowledge no previous studies have assessed the effects of prolonged cyclic loading on ALP activity. Here, we found that prolonged cyclic loading yielded a return to baseline in the middle and transition regions and the formation of a similar ALP gradient to slow stretch, indicating that growth rate may be responsible for early initiation of mineralization and cyclic loading may affect later stage mineralization.

Taken together, this data suggests the formation of a fibrocartilage-like tissue gradient mirroring native enthesis development. Early bone development is characterized by the presence of types II and X collagen, as well as aggrecan [3]. Upon further maturation, the localization of these key enthesis proteins shifts to the developing enthesis interface ultimately giving rise to the compositional gradients underpinning mature fibrocartilage and mineralized fibrocartilage [9,10]. Previously, we found 6 weeks of static clamping increased GAG accumulation and ALP activity, as well as type II collagen, type X collagen, and aggrecan localization under the clamp, mirroring early enthesis development via endochondral ossification [12]. In this study, the addition of slow stretch and cyclic loading yielded similar trends in GAG accumulation and ALP activity at an earlier timescale, and differentially drove a shift in type II collagen, type X collagen, and aggrecan to the transition and clamped regions. Collectively, this further suggests that slow stretch and cyclic loading differentially contribute to the development of more mature, late postnatal-like engineered entheses.

A key finding of this study was that slow stretch and cyclic load differentially drove improvements in zonal collagen organization. Interestingly, only slow stretch yielded a shift in the transition region from more perpendicular to more parallel, direct insertions, similar to the shift in collagen organization in developing entheses [2,3,20]. To better understand if this shift was dependent on the load type or if additional cyclic loading could drive a similar shift in organization, we compared 10 days versus 20 days of cyclic load (24 days versus 34 days of total culture). We found no further maturation in the transition region with an additional 10 days of cyclic loading. This further suggests that the shift in organization to more direct insertions is load rate dependent and driven by slow stretch. Mirroring the lack of organizational improvement in collagen organization, prolonged cyclic stretch did not improve mechanical properties and had little impact on zonal composition and protein localization, except for a shift in zonal ALP activity. Collectively, this suggests that slow stretch and cyclic load differentially drive maturation and indicates the need for combined or ramped loading to further drive maturation at the insertion site in tissue engineered constructs.

By using an *in vitro* culture system that mirrors enthesis postnatal development we investigated the individual effects of mechanical cues that mimic ACL growth rate and cyclic muscle activity on ligament-to-bone enthesis development and maturation. This study demonstrated that slow stretch and intermittent cyclic load differentially drive maturation of zonal collagen organization and accumulation of types II and X collagen, both resulting in improved transition region mechanics. Notably, slow stretch resulted in more direct fibril and fiber insertions in the transition region, mirroring late postnatal enthesis maturation, whereas cyclic load resulted in improved fiber and fascicle organization in the middle region, mirroring ligament maturation. Additionally, this study demonstrated that prolonged cyclic loading did not result in further improvements or a shift in collagen organization in the transition, indicating that slow growth elongation could be the driver of this shift in enthesis development. These results are promising for better understanding enthesis development and for better informing how to leverage developmental mechanical cues to engineer bone-ligament-bone replacements; however, this study is not without limitations. Slow stretch and cyclic loading yielded promising results demonstrating the possibility that growth rate and muscle loading may differentially drive local enthesis maturation; however, only one loading regime for each respective mechanical cue was analyzed in this study. Of note, embryonic ACL growth rate of 1 mm/day was also analyzed [37]; however, this resulted in a 100% failure rate by 10 days with constructs failing as early as 3 days after initiation of load, indicating the possible need for earlier loading or phased loading to better mirror development. Additionally, cyclic loading was not found to drive improvements in organization at the interface even with prolonged loading. The loading regime used has been shown to be optimal for anabolic and catabolic response in engineered constructs [39,58,59], and it is within the range of physiological human ACL strain (2-8%) [60–63]. However, additional cyclic strain rates, amplitudes of strain, or durations of loading may yield improved organizational maturation at the interface compared to 5% strain. Future work should also investigate the synergistic effects of slow stretch and cyclic loading to further drive accelerated development of native-like engineered entheses.

Despite these limitations, the ligament-to-bone entheses developed in this study are some of the most physiologically relevant engineered entheses to date. Additionally, by leveraging an *in vitro* culture system that mimics enthesis development, developmental mechanical cues that are difficult to differentiate *in vivo* were easily separated and investigated. This provides a promising tool that can be further used to rapidly and cost effectively investigate additional mechanical loading regimes, possibly leading to improved rehabilitation protocols, engineered replacement options, and regeneration of the enthesis following graft repair.

## 5. Conclusions

This study provides new insight into the differential effects of slow stretch and cyclic loading in tissue engineered ligament-to-bone entheses. It is known that enthesis postnatal maturation is largely driven by mechanical cues; however, it has been challenging to separate these cues *in vivo* to evaluate the differential effects on enthesis maturation, limiting repair options. Notably, we have demonstrated that slow stretch drives maturation of the enthesis/transition region, whereas cyclic load drives maturation in the ligament mid-substance. These constructs are some of the most organized bone-ligament-bone constructs developed to date and are promising as future functional ACL replacements. Further, the *in vitro* system used in this study closely mirrors enthesis development and is a promising tool to quickly investigate and understand how different mechanical cues can drive enthesis maturation or regeneration leading to improved engineered replacement options or rehabilitation protocols following injury.

## Acknowledgements

The authors would like to thank Dr. Ning Zhang and Anne Katherine Brooks for their assistance with this study. The authors would like to acknowledge the use of facilities and instruments within the Nanomaterials Characterization Core at Virginia Commonwealth University. The authors acknowledge that services and products in support of this study were generated by the Virginia Commonwealth University Cancer Mouse Models Core Laboratory, supported in part with funding from NIH-NCI Cancer Center Support Grant P30 CA016059. This study was supported, in part, by a grant from the Orthoregeneration Network (ON) Foundation, Switzerland (19-037), Koerner Foundation Fellowship, and PI Startup Funds.

## Declaration of competing interest

The authors declare that they have no known competing financial interest or personal relationships that could have appeared to influence the work reported in this paper.

## Data Availability

The raw/processed data required to reproduce these findings cannot be shared at this time as the data also forms part of an ongoing study. However, research data is available upon request to the corresponding author.

## Notes

### Competing Interest Statement

The authors have declared no competing interest.

