## Supplemental for "Enthesis Maturation in Engineered Ligaments is Differentially Driven by Loads that Mimic Slow Growth Elongation and Rapid Cyclic Muscle Movement"

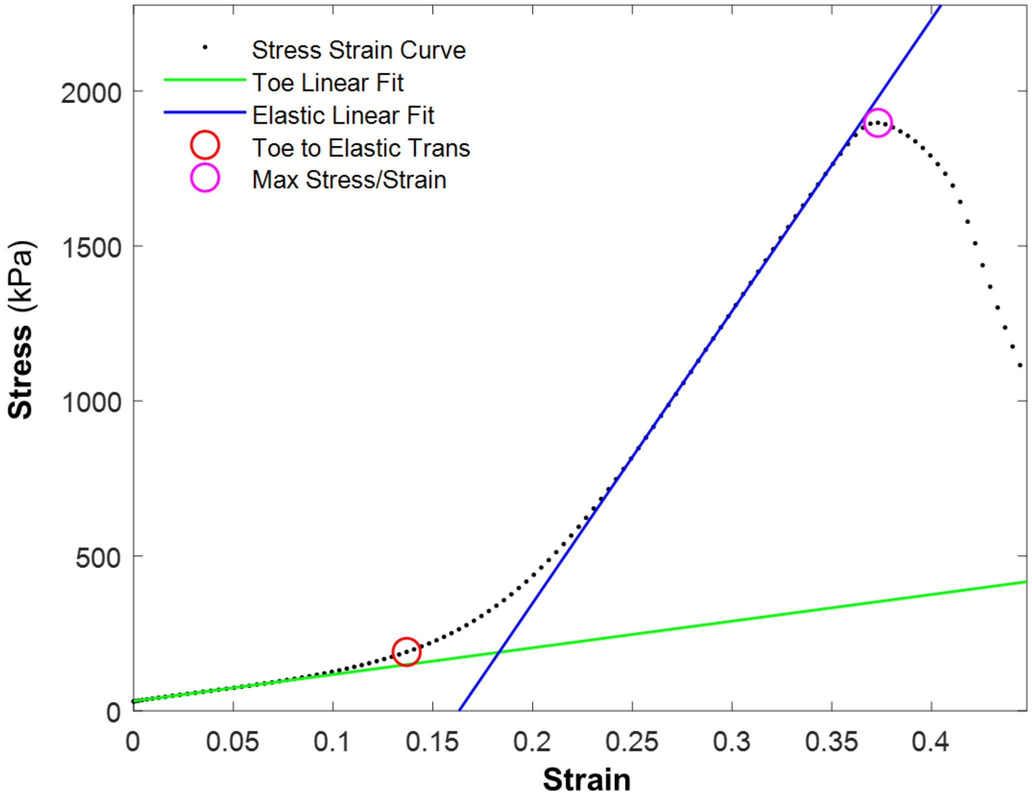


**Supplemental Figure 1:** Example graphical output of Matlab code for calculation of mechanical properties. Properties include stiffness and elastic modulus (depicted by blue line for elastic linear region fit), toe modulus (depicted by green line for toe linear region fit), ultimate tensile strength and strain at failure (depicted by pink circle), transition stress and strain (depicted by red circle indicating shift from toe region to elastic region).

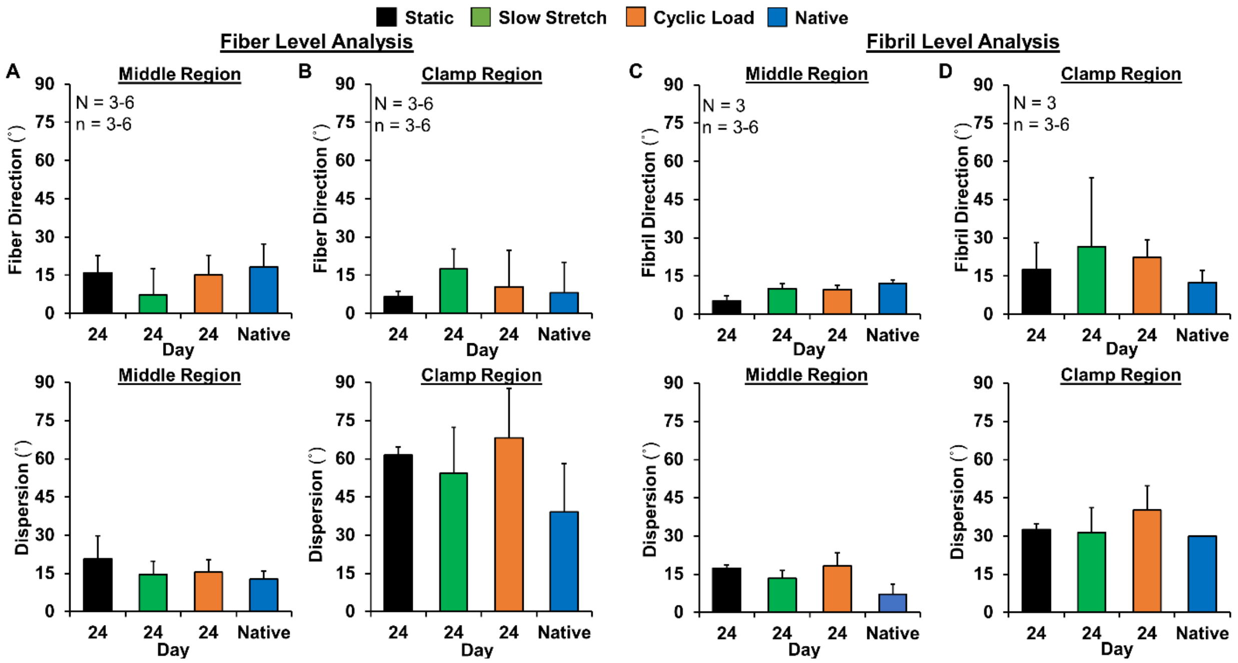


**Supplemental Figure 2:** Fiber level image analysis of confocal reflectance images of (A) middle and (B) clamped tissue regions at 24 days compared to native immature bovine ligament and bone, respectively. Fibril level image analysis of SEM images of (C) middle and (D) clamped tissue regions at 24 days compared to native immature bovine ligament and bone, respectively. Both fiber and fibril angle mirrored immature bovine ligament in the middle region with minimal dispersion. Organization mirrored native bone under the clamp with large dispersion values, indicating a lack of directional organization. 3-6 images were averaged per region for each construct or native tissue sample to determine construct averages. The reported angle of alignment and dispersion are the average and standard deviation of the construct averages for each tissue region (N=6 for engineered tissues in A &B, N=3 for engineered tissues in C&D and N=3 for native tissue). Significance compared to *bracket ($p < 0.05$).

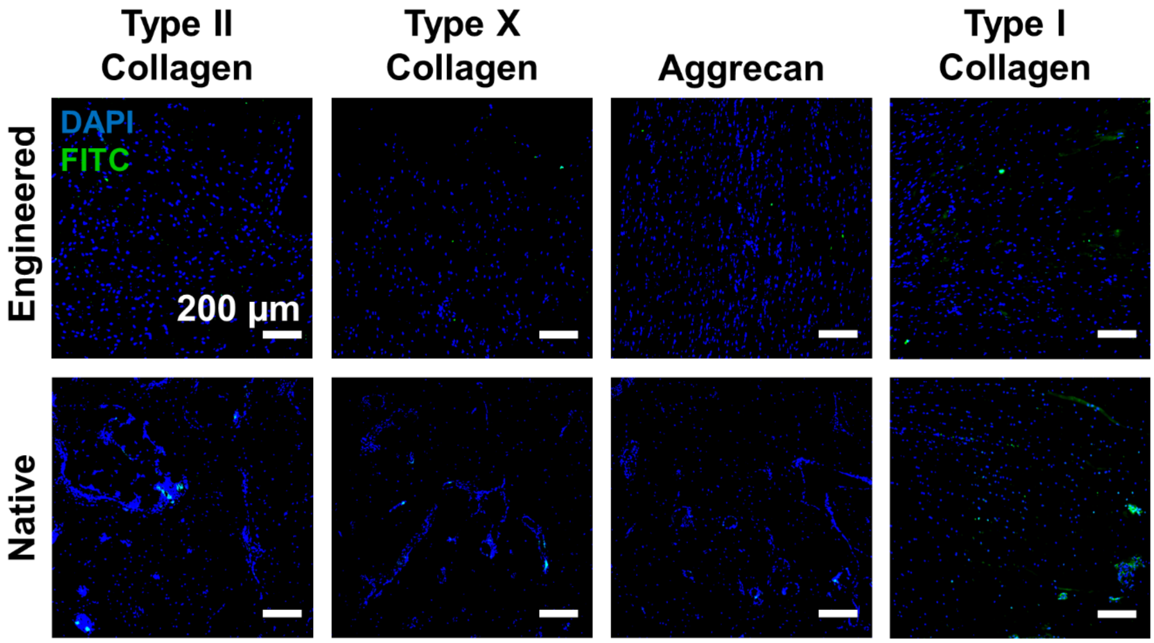


**Supplemental Figure 3:** Negative immunofluorescence images of the transition region of tissue engineered constructs at 24 days and of enthesis tissue regions of native samples are shown for type II and X collagen, aggrecan, and type I collagen. FITC = representative protein, DAPI = nuclei, scale bars = 200 µm.
